# A Key For Hypoxia Genetic Adaptation In Alpaca Could Be A HIF1A Truncated bHLH Protein Domain

**DOI:** 10.1101/386987

**Authors:** M Daniel Moraga, Fernando A. Moraga C, Felipe Figueroa

## Abstract

Animals exposed to hypoxia, triggers a physiological response via Hypoxia Inducible Factors (HIF1). In this study, we have evidenced the existence of genetic events that caused the loss of most of the bHLH domain in HIF1A proteins borne by Alpaca and other members of the *Cetartiodactyla* superorder. In these truncate domains, some stop codons are found at identical nucleotide positions in both, Artiodactyls and Cetaceans, indicating that mutations originating the truncated domains occurs before their divergence about 55 million years ago. The relevance of this findings for adaptation of Alpacas to hypoxia of high altitude conditions are discussed.

## Introduction

Camelids and remaining even-toed ungulates (artiodactyls) together with whales and dolphins (cetaceans) are grouped in the superorder *Cetartiodactyla,* Price *et al*. (2005). Alpacas (*Lama pacos* Linnaeus, 1758), reclassified as *Vicugna pacos* by Kadwell *et al.* (2001), is one of the four species of South American camelids. Llamas (*Lama glama* L) along with Alpacas are domestic species, while guanacos (*Lama guanicoe* Miller 1776) and vicuñas (*Vicugna vicugna* Molina 1782) are wild species.

During the evolution of South American camelids, they developed physiological adaptations to the cold environments and food shortages typical of environments of high-altitude hypoxia, 3000 meters over the sea level Wheeler (2012). Hypoxia is a situation in which there is a reduction in the availability of oxygen to tissues and cells. Different physiological adaptations for living in hypoxia has been described in Llamas. These, include higher affinity of hemoglobin for O_2_, slight increase in blood hemoglobin concentration, high muscle myoglobin concentration, a more efficient O_2_ extraction at tissue levels, high lactic dehydrogenase activity, and less muscularized pulmonary arteries, Llanos *et al.* (2011a, 2011b).

Hypoxia is associated with developmental, physiological environmental and pathophysiological conditions as ischemia, arthritis, inflammation, chronic lung disease, stroke, heart disease and cancer, Semenza (1999). A particular archetype of hypoxia is that associated with high altitudes (hypobaric hypoxia). As a consequence of this type of hypoxia, humans and animals triggers an acute (AMS) or chronic (CMS) response depending on the time of exposure. AMS is established when a person is exposed for a short period to hypobaric hypoxia and develops signs of a headache, fatigue, sleep disorder, gastrointestinal disorders or vertigo, Davis *et al.* (2017). CMS, also known as Monge’s disease, may be come about when a person lives for a long time at high altitude, reviewed by Villafuerte & Corante (2016). Around 1.2 to 33% of populations living at high altitude suffer from CMS depending on factors such as age, sex, high and the origin of the population, Azad *et al.* (2017). One important sign of CMS is the elevation of the hematocrit and the number of erythrocytes (polycythemia). While increasing the amount of hemoglobin in the blood could be a beneficial adaptation to hypoxia, excessive erythrocytosis results in a higher blood viscosity, which affects tissue blood flow and oxygen supply Prchal (2010).

The Hypoxia Inducible Factors (HIFs) proteins are transcription factors that play a key role in the homeostasis of oxygen in animals, Rytkonen & Storz (2011). HIFs proteins are basic Helix-Loop-Helix–PER–ARNT–SIM (bHLH–PAS) proteins distinguish by being both ubiquitously expressed and signal-regulated or constitutively active and tissue specific, Button *et al.* (2017). HIFs proteins form heterodimeric complexes that are composed of alpha O_2_-labile subunit, either HIF1A, EPAS1 (HIF2A α) or HIF3A and one stable β-subunit HIF1β also known as Aryl hydrocarbon receptor nuclear translocator (ARNT). In normoxia conditions in presence of O_2_, the HIFA subunits are modified by the HIF-specific prolyl-hydroxylases (PHD), causing proteasomal degradation mediated in part by the von Hippel-Lindau suppressor protein (VHL), Button *et al.* (2017), Fribourgh & Partch (2017).

Ethiopians in Africa, Han in Asia and Andean populations of South America have been the most investigated to elucidate the adaptive mechanisms to CMS of high altitude human populations, Villafuerte & Corante (2016). Several of these studies have demonstrated the existence of a genetic adaptation to hypoxia in Tibetan populations, for review see Murray et al. (2018). Genetic adaptation to hypoxia conditions has also been evidenced in goat, dog, and sheep that live in high-altitudes, Song *et al.* (2016); Wang *et al.* (2014); Zhang *et al.* (2014).

In the present study, we have investigated the phylogenetic relationships and genetic structures of the HIF1A proteins carried by members of the superorder *Cetartiodactyla*. During our investigation, we discovered the existence of a genetic event that caused the loss of most of the bHLH domain in the proteins borne by the Alpaca and other members of the *Cetartiodactyla* superorder. As the mutations affect, both the Artiodactyls and Cetaceans, we postulate that the mutation occurred before their divergence about 55 million years ago, Gingerich & Uhen (1998). The relevance of these findings for genetic adaptation of Alpacas to hypobaric hypoxia of high altitude conditions is discussed.

## Material and methods

### The species dataset

The complete set of HIF1A protein sequences of the superorder *Cetartiodactyla* included in the present study belong to 38 species of families from orders *Artiodactyla* and *Cetacea*. The species have been identified by a 4-digit acronym, in which the first two letters correspond to the name of the genus and the last two, to the name of the species. These acronyms will be used from here on.

The families of the Order *Artiodactyla* are *Camelidae*: *Vicugna pacos* (Vipa); *Camelus bactrianus* (Caba); *Camelus dromedarius* (Cadr) and *Camelus ferus* (Cafe). *Bovidae*: *Bison bison* (Bibi); *Bos taurus* (Bota); *Bos mutus* (Bomu); *Ovis aries* (Ovar); *Pantholops hodgsonii* (Paho); *Capra hircus* (Cahi) and *Bos grunniens* (Bogr). *Cervidae*: *Odocoileus virginianus texanus* (Odvi). *Suidae*: *Sus scrofa* (Susc).

The families of order *Cetacea* are *Balaenopteridae*: *Balaenoptera acutorostrata scammoni* (Baac). *Phocoenidae*: *Neophocaena asiaeorientalis* (Neas). *Physeteridae*: *Physeter catodon* (Phca). *Delphinidae*: *Orcinus orca* (Oror) and Tursiops *truncates* (Tutr). *Monodontidae*: Delphinapterus *leucas* (Dele). *Lipotidae*: *Lipotes vexillifer* (Live).

### Selection of HIF1A proteins

Proteins containing PAS domains borne by Cetartiodactyls species were retrieved from the NCBI GenBank (www.ncbi.nlm.nih.gov). We used a PSI-BLAST search with three iterations and the Alpaca EPAS1 protein (accession number XP_015105262.1) and the sequences were subsequently grouped with the help of the CLANS program, Zimmermann *et al.* (2017). Later, only HIF1A proteins were selected and aligned with the help of the Clustal Omega program and further analyzed.

### Dendrogram construction

Evolutionary relationships were evaluated by genetic distance methods using the neighbor-joining algorithm for phylogenetic tree construction of the Mega 7 program, Kumar *et al.* (2016). The parameters pairwise deletion and p-distance model were used. Bootstrap test of phylogeny was performed with 1000 replicates.

## Results

As the EPAS1 (HIF2A) protein has been implicated in the genetic adaptation to the hypoxia conditions in humans and animals living in the plateau altitudes, Murray *et al.* (2018); Song *et al.* (2016); Wang *et al.* (2014); Zhang *et al.* (2014), we first searched in the *Vicugna pacos* NCBI database for corresponding EPAS1 protein sequences. A single sequence was found (accession number XP_015105262.1) which then we use as a query, to perform a BLAST search for PAS-containing proteins on the whole *Cetartiodactyla* NCBI DataBank. A total of 959 sequences were identified and retrieved, 46 of which correspond to Alpacas. In the present study, we restricted the genetic analysis to the 38 HIF1A proteins. Remaining analysis of others PAS proteins will be published somewhere else.

The set of the 38 HIF1A proteins sequences included in the present study are, from the order *Artiodactyla*, (1 Vipa sequence; 1 Caba; 2 Cafe; 1 Cadr; 2 Bibi; 2 Bota; 2 Bomu; 2 Ovar; 3 Paho; 4 Cahi; 3 Bogr; 2 Odvi; 1 Susc) and from the order Cetacea (2 Baac; 1 Neas; 3 Phca; 1 Oror; 1 Tutr; 1 Dele; 3 Live).

The alignment of the sequences is shown in Figure 1 and in the Supplementary A section. In Figure 1 the alignment include the first 480 amino acids of 12 representative HIF1A protein sequences encompassing the bHLH, PAS and PAS 3 domains. In the Supplementary A section, the whole set of the 38 full length HIF1A proteins is included. The alignment of the protein sequences reveals that they can be divided according to their length, into two categories which we designate as Long and Short (L or S in Figure 1 and Supplementary A section). The former start with a Met amino acid at position 1, 4 or 14 of the alignment, the latter with a Met amino acid at position 63. Of the all sequences, 28 belong to the L type, while the remaining 10 correspond to the S type. Sequences of L or S type are present in both, *Artiodactyla* and *Cetacea* orders. For example, among the S type sequences are those borne by the Artiodactyls Vipa, Cafe, Cadr, Cahi and Bibi and by the Cetacean Baac, Tutr, Phca, Odvi, and Live. A similar situation occurs with the L type sequences (Figure 1). When in the alignment more than 1 HIF1A proteins are borne by members of one single species, the sequences may be either of the same types or from a different one. Examples from the first category are the 4 Paho sequences which all belong to the L type. Alternatively, out of the four Cahi sequences, 3 are L and 1 S, or the Phca sequences (2 L and 1 S) and Baac sequences (1L and 1 S).

**Fig 1.**
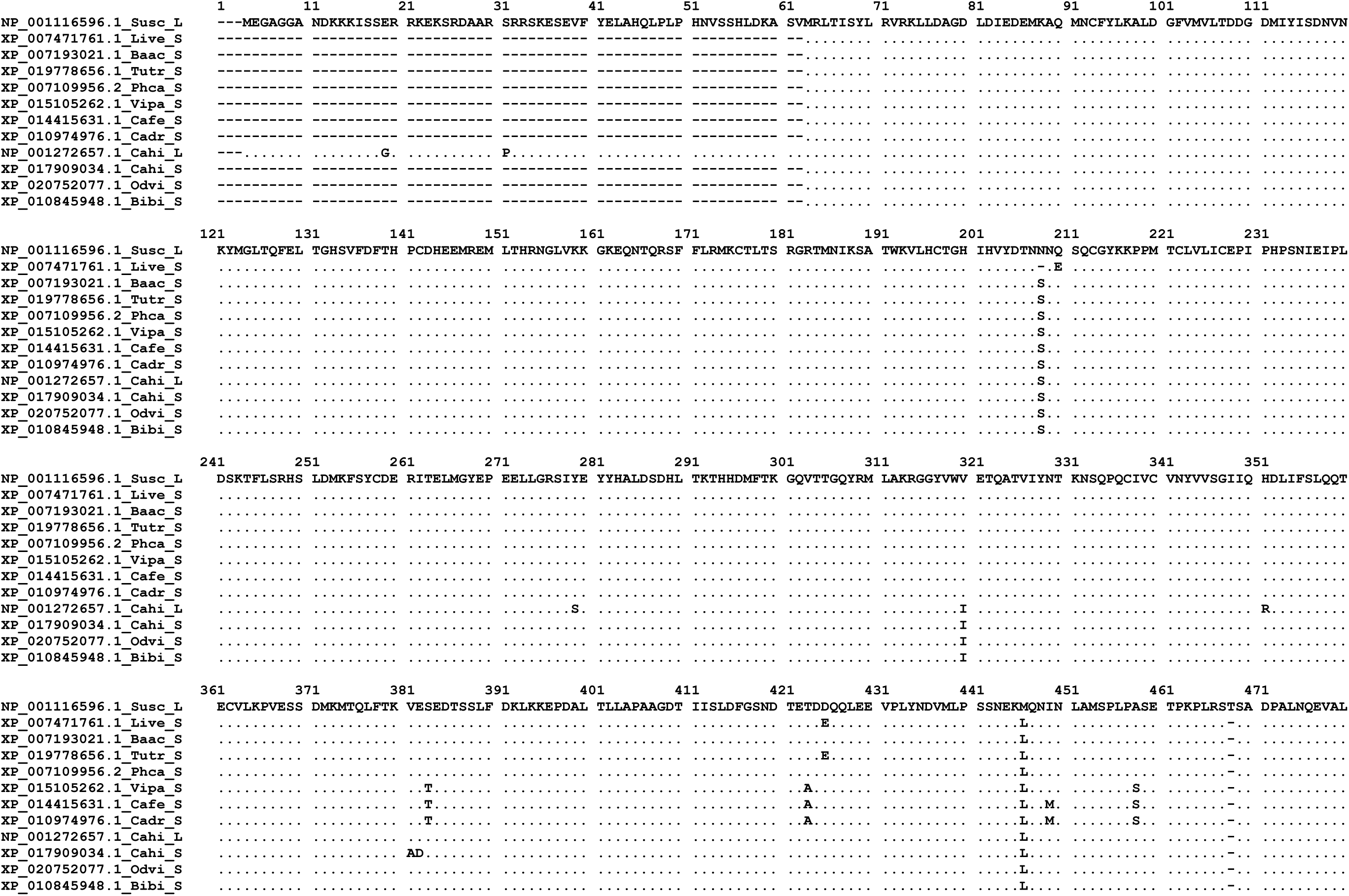
Alignment of *Cetartiodactyla* HIF1A proteins. Dots indicate identity with the sequence of the top of the figure. Dashes from site 1 to position 62 in short sequences indicate deleted amino acids in the bHLH domain of the HIF1A proteins. Dashes at the end of the proteins indicate unavailable information. Name of the species after accession numbers corresponds to the mention in the material and methods section.

In a characteristic HIF1A protein, four domains are present. A basic Helix-Loop-Helix amino-terminal domain (bHLH), is followed by two PER-ARNT-SIM domains (PAS) and then by two transactivation domains, one NH2-terminal domain and one COOH-terminal domain. In the set of sequences that we have analyzed, we have realized that the short sequences differ from the long ones, in that they are devoid of most of the bHLH domain. In fact, only the second helix region is present. In the long type, the entire bHLH domain contained 50 amino acids, while the short type contains only 13 residues. Thus, in the short, only amino acids corresponding to the second helix region are present (numbering according to Zhou *et al.* 1997).

Sequence analysis at the DNA level allows getting an insight into the genetic events that generated the short HIF1A proteins. The alignment of the sequences is shown in Figure 2 and in the Supplementary B section. In Figure 2 the alignment include the full bHLH domain of 15 representative HIF1A protein sequences. In the Supplementary B section, the whole set of the 34 full length bHLH domain sequences is included. In the alignment, the conserved sequence CGAAAAGAG in position 37-39, encodes for the first 3 residues (RKE in Figure 1), of the bHLH domain of HIF1A proteins. Although the next 5´upstream 16 positions are also well preserved, then the sequences are segregated into two clusters. One of the clusters, in which all short sequences are grouped, is distinguish from the other group in having a high nucleotide variability and been very Thymine rich at its 5´end.

**Fig 2.**
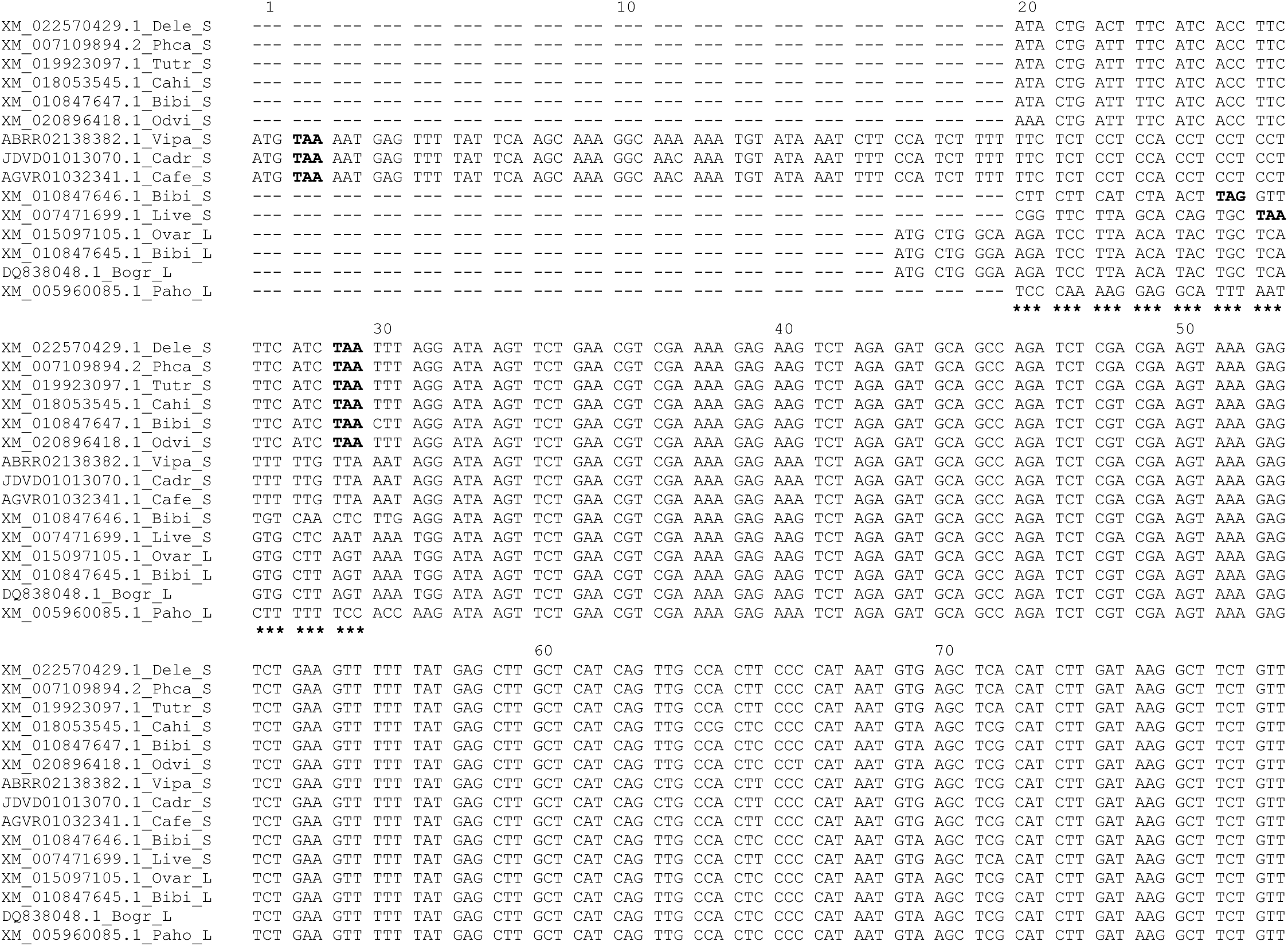
Nucleotide alignment of *Cetartiodactyla* bHLH domain of HIF1A proteins. Dashes at the beginning of the sequences were introduced for optimal alignment. Asterisk at the bottom of sequences between position sites 20-29 indicate the Thymine rich stretch of the sequences. Stop codons are in bold letters. Name of the species after accession numbers corresponds to that mention in the material and methods section.

In the alignment, the Cetacean sequences Dele, Phca, Tutr, Odvi and Baac and the Artiodactyls Cahi and Bibi conform an identical or nearly identical compact group and in all seven sequences a TAA stop codon is present at position 29 of the alignment. All these sequences are short and the presence of the stop codon by itself explain the fact of the shortening of the HIF1A protein that they borne. Sequence Live, which also have de TAA stop codon in the Thymine rich stretch is different from all other short nucleotide sequences. Finally, camelids Vipa, Cadr and Cafe conform a second conserve cluster, with a stop codon TAA at position 2 in the alignment.

## Discussion

When animals are acute or chronically exposed to hypobaric hypoxia, they trigger a hypoxic response via Hypoxia Inducible Factor (HIF) proteins that function as transcriptional complexes. HIF proteins are heterodimers with 2 chains, HIF-alfa and HIF-beta. A full hypoxic response requires dimerization, nuclear translocation and binding of HIF proteins to p300-CBP proteins, important for maximal transcriptional activation, Arany *et al.* (1996).

The analysis of the sequences HIF1A proteins in *Cetartiodactyla* reveals that the Alpaca protein lacks most of the bHLH domain, present in the generality of bHLH-PAS family proteins, Wang *et al.* (2014). These proteins share a common structure that includes DNA Binding domain (bHLH), followed in tandem by PAS domains (PAS-A and PAS-B) and variable domains of transactivation or transrepression.

The alignment in Figure 2 provides several distinctive features which give some insight into the evolution of the nucleotide bHLH encoding domain. First, all but one of the seven sequences at the top part of the Figure 2 are identical in 5´thymine rich stretch. The only exception is the Odvi sequence. Second, all the seven sequences contain the stop codon TAA at position 29 of the alignment. Third, in the Neighbor joining (NJ) tree shown in Figure 3, the seven sequences are grouped into two branches. One contains the three artiodactyl sequences (Cahi, Odvi and Bibi) and the second all the four cetacean bHLH sequences. As a whole, these three characteristics seem to indicate that in the evolution of HIF1A protein, the advent of the short protein version was a unique mutational event which gave rise to a bHLH pseudogene. Also, the data indicate that this mutational event occurred before the divergence of artiodactyls and cetaceans, about 55 Mya Gingerich & Uhen (1998).

**Fig 3.**
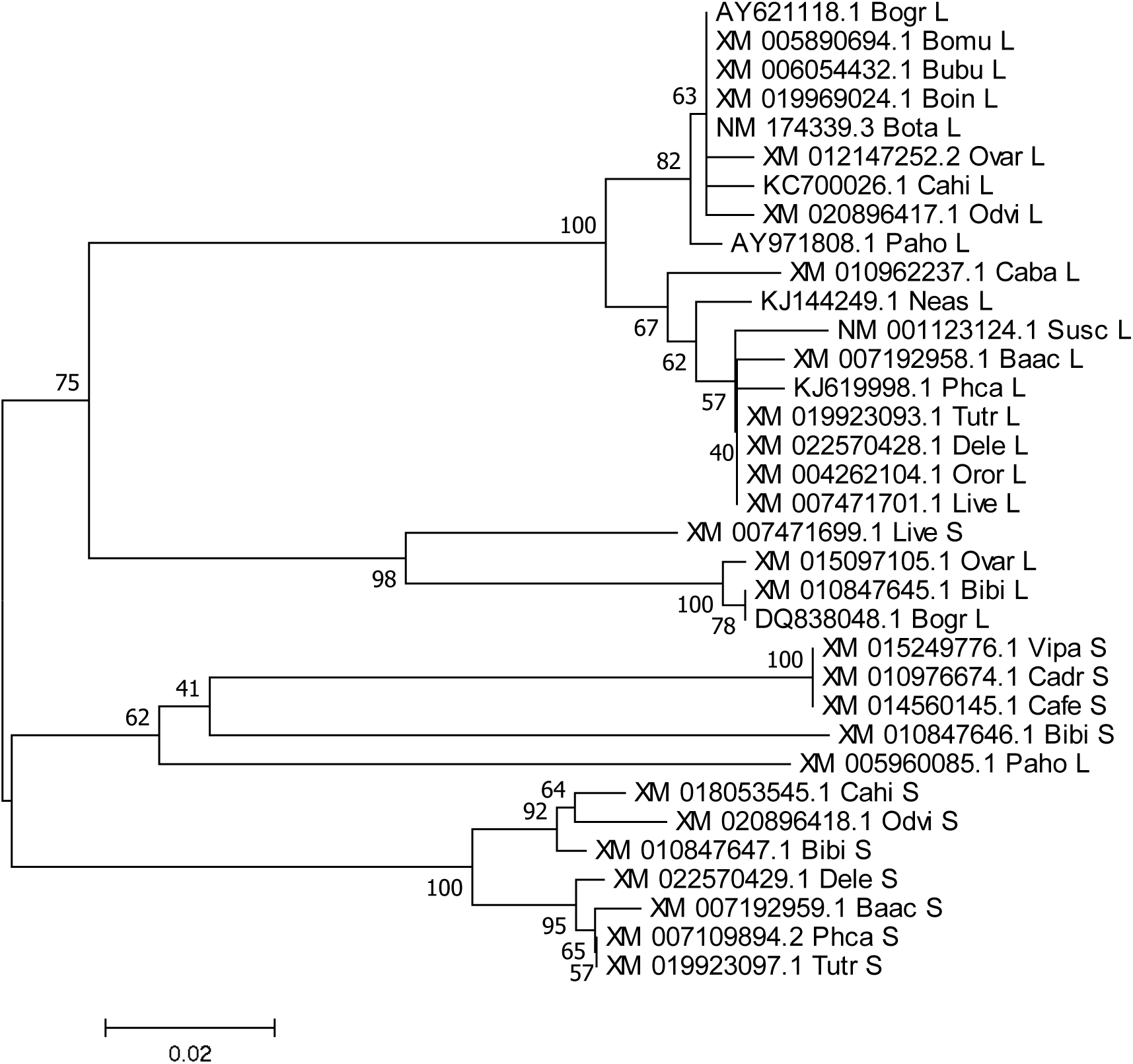
Phylogenetic trees based on selected nucleotide bHLH domain of *Cetartiodactyla* HIF1A sequences. The numbers shown on the interior nodes are bootstrap probabilities in percent. The parameters pairwise deletion and p-distance model were used.

Additional information is provided by the camelid species, one Alpaca (Vipa) and the two camel sequences (Cadr and Cafe), which all three share the 5´thymine richness nucleotide stretch with the other short sequences. The three sequences also show a TAA stop codon, but this is located 27 codons upstream of one detected in the previous seven ones. All three camelids sequences are almost identical each other which indicate that their differentiation with the rest of the sequences occur during the 40-45 Millions year of evolution in the plains of North America and before the separation of North and South American camelids, which occurs some 3 Mya, Wheeler, 2012). The short Bibi sequence is also 5´thymine rich and also contain a stop codon, but this is instead TAG and is located 4 codons upstream of the TAA in sequences at the top of Figure 2. Finally, the S type Live sequence is similar to all other short sequences in having both the TAA stop codon and in being Thymine rich at its 5´end. A particular feature of the sequence is the fact that it shows at the 5´end some similarities with the three of the long sequences Ovar, Bibi and Bogr. Taking this similarity into account, we can speculate that perhaps the original mutation originating the bHLH pseudogene could have occurred in a long sequence which was rich in thymine at its 5´end.

A phylogenetic tree based on bHLH nucleotide sequences shows that they are grouped with two exceptions, into two major clades depending on whether they are derived from coding sequences for long or short HIF1A proteins (Figure 3). In one of the clades are found all but one of the long sequences. In the other clade are found all but one of the short sequences. The exceptions are L Paho and the S Live sequences, which incongruous locations can be explained by their differences at a stretch from codon 20 to 30 in the alignment.

In the clade of the long sequences, three different branches can be found. In the first are all Bovidae sequences, such as Bota and Ovar and one of the two sequences of Paho. The second branch contains the long sequences derived from the *Camelidae* Caba, the *Cervidae* Neas, the *Suidae* Susc and all the long sequences derived from cetaceans. In a third branch are the 5´Thymine rich long *Bovidae* Ovar, Bibi and Bogr sequences and the incongruous *Lipotidae* Live short sequence. In the clade of the short sequences, two main branches can be found. One contains the Camelids, Cadr, Cafe and Vipa, the Bovids Bibi and the incongruous Paho long sequence. The others are all the seven sequences described above.

Initial studies by Reisz-Porszasz *et al.* (1994) showed that cutting the bHLH domain does not affect the dimerization and nuclear translocation of HIF1A, but it diminished its binding to DNA, impelling the normal hypoxic response. Further studies have corroborated these conclusions (Semenza *et al.* 1997; McGuire *et al.* 2001; Huang *et al.* 2012). Then, according to these findings, we conclude that the hypoxic-response in Alpacas is diminished due to the adaptive loss of part of bHLH domain of HIF1A proteins. Consequently, Alpacas are genetically adapted to live in a low oxygen tension environment, without the negative consequences of a full hypoxic response, such as AMS and CMS. Interestingly, we have found that the absence of most the bHLH domain in the HIF1A protein of the only South American camelid included in the present study, also seems to exist in other members of the *Cetartiodactyla* superorder. Thus, in addition to the Alpaca, truncated bHLH domain in HIF1A proteins, are also possible to be found among goats, old world camelids and cetaceans. The main conclusion of this finding is that the mutation(s) that gave rise to truncated bHLH domains in some *Cetartiodactyla* HIF1A proteins, occurred around 55 Mya, before the divergence of *Artiodactyla* and *Cetaceans* Gingerich & Uhen (1998).

## Acknowledgments

We thank Mr. Ricardo Reyes for the help in editing the manuscript. There are none funding sources.

## Supplementary material A

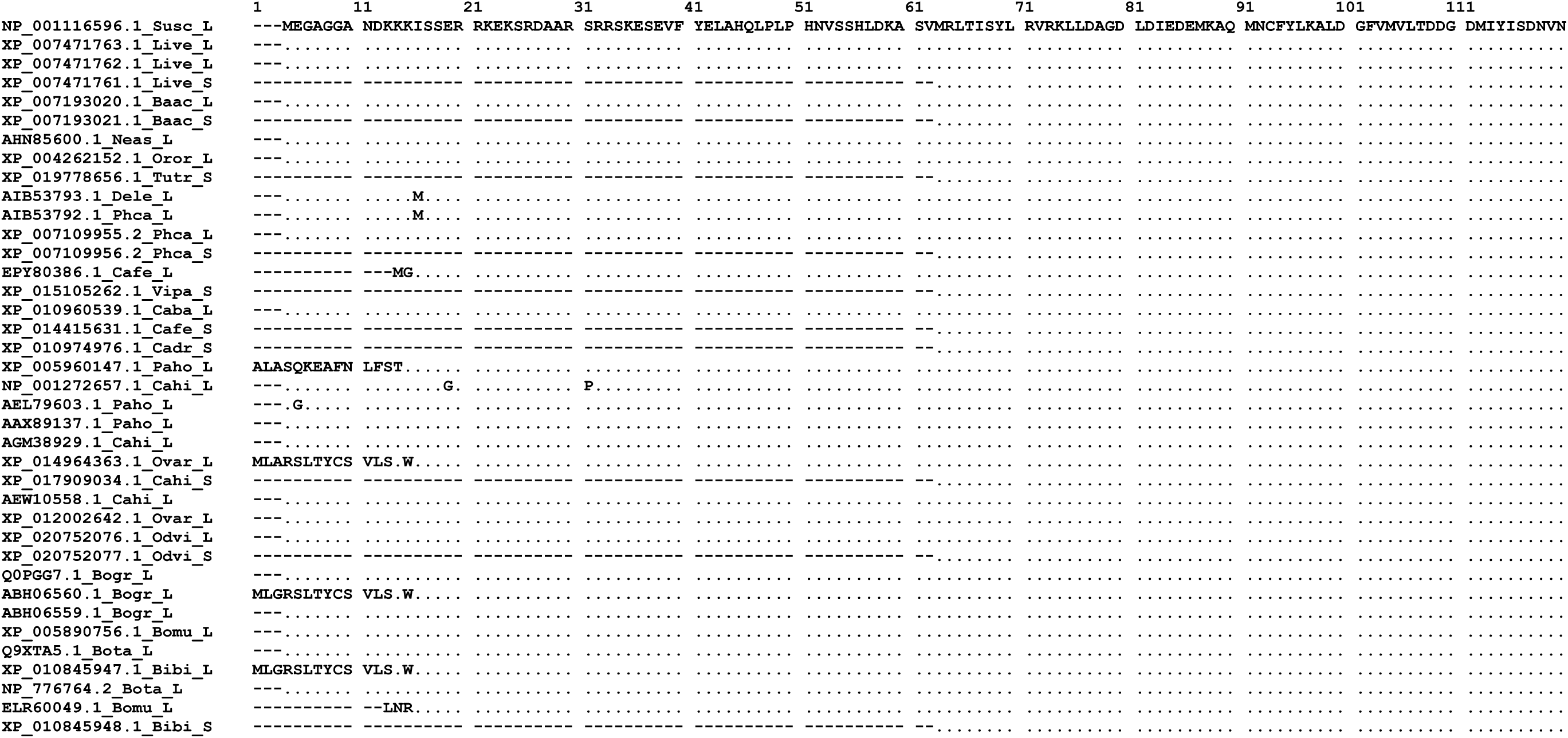
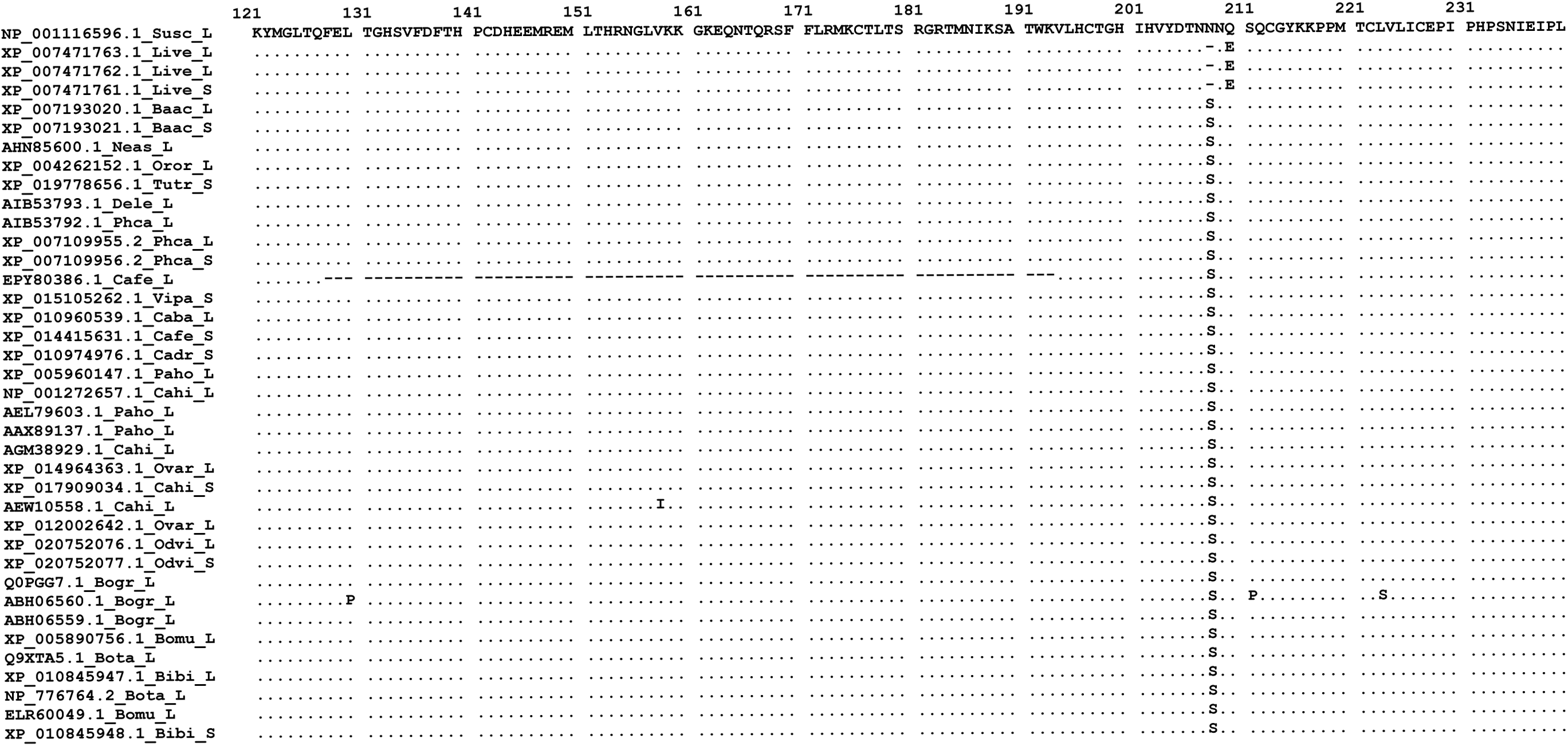
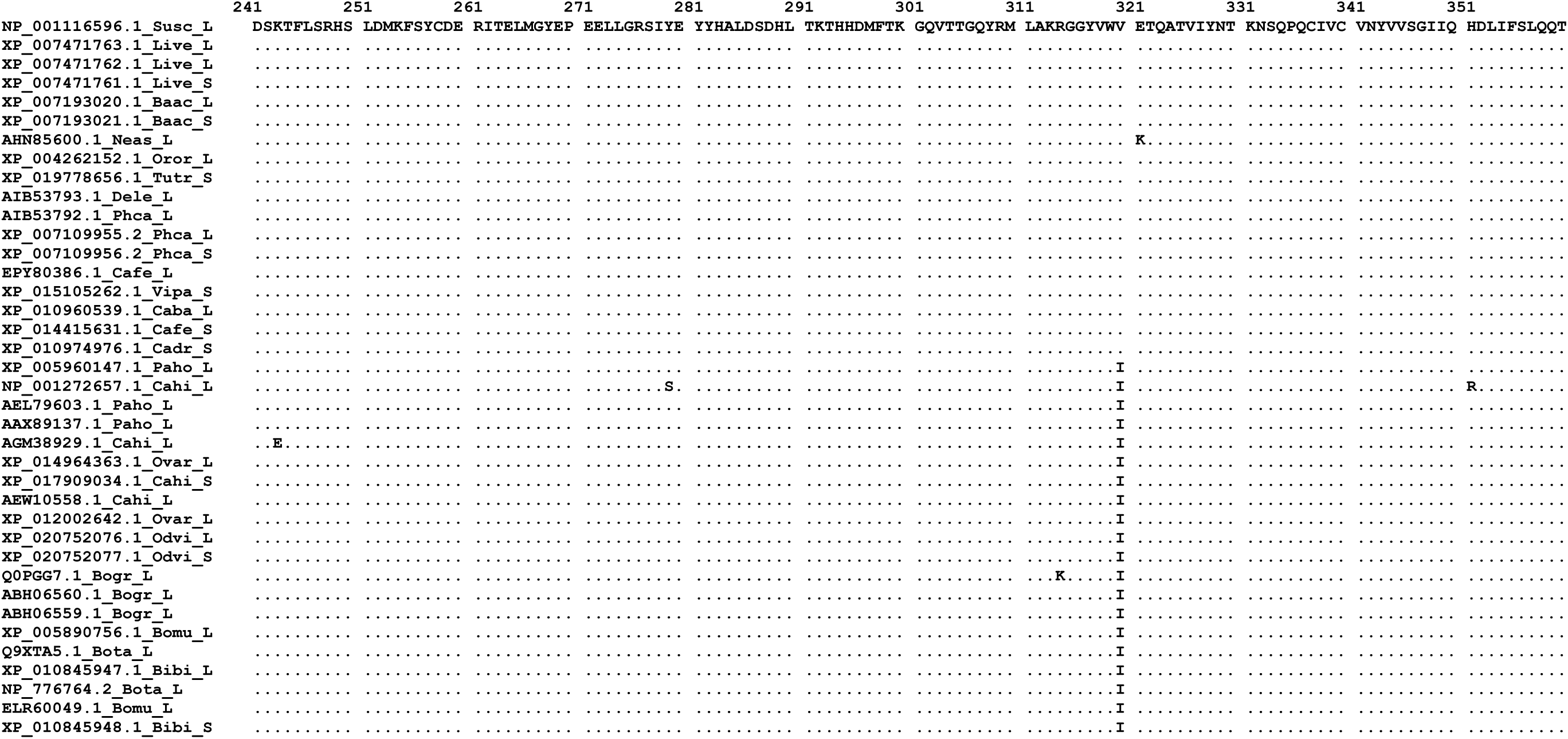
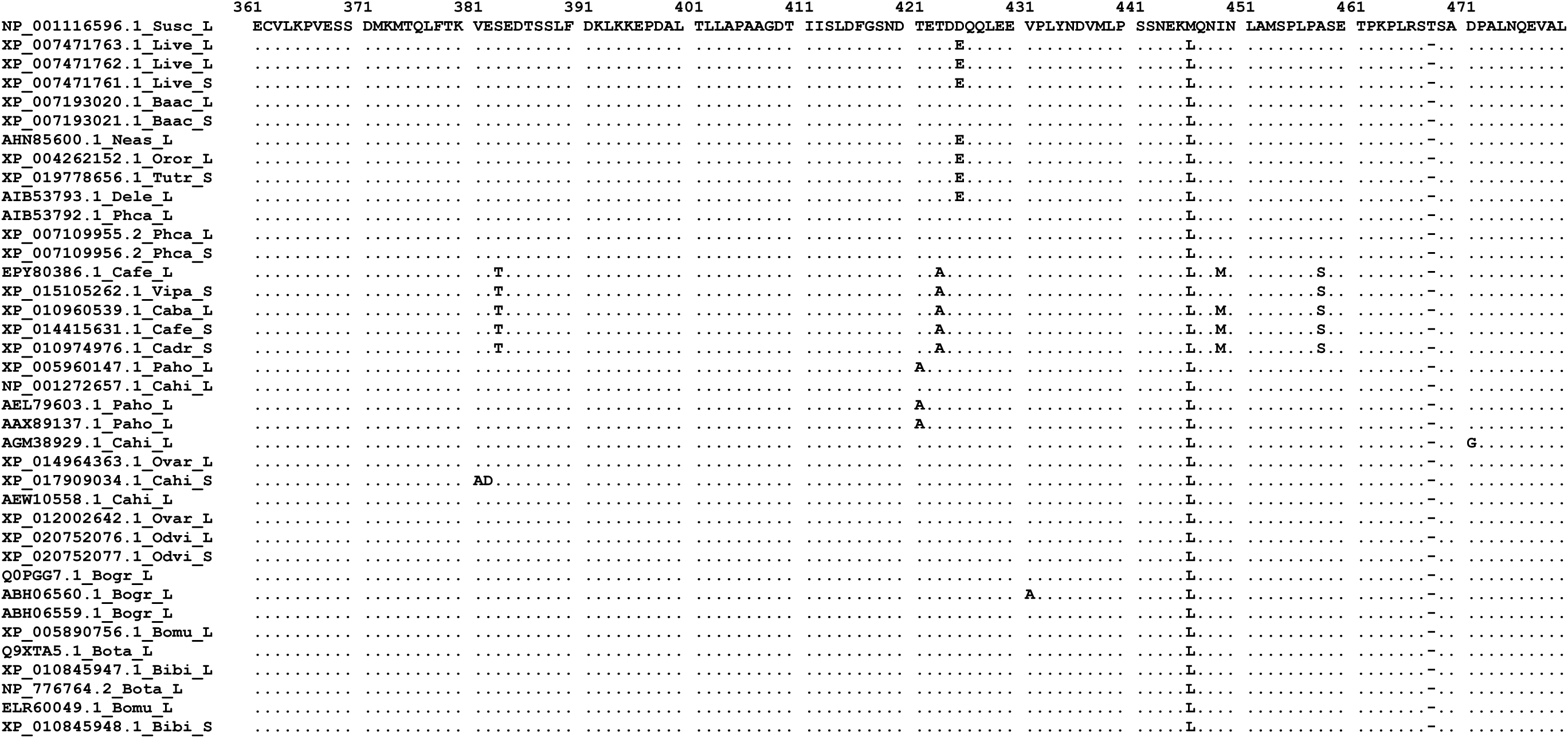
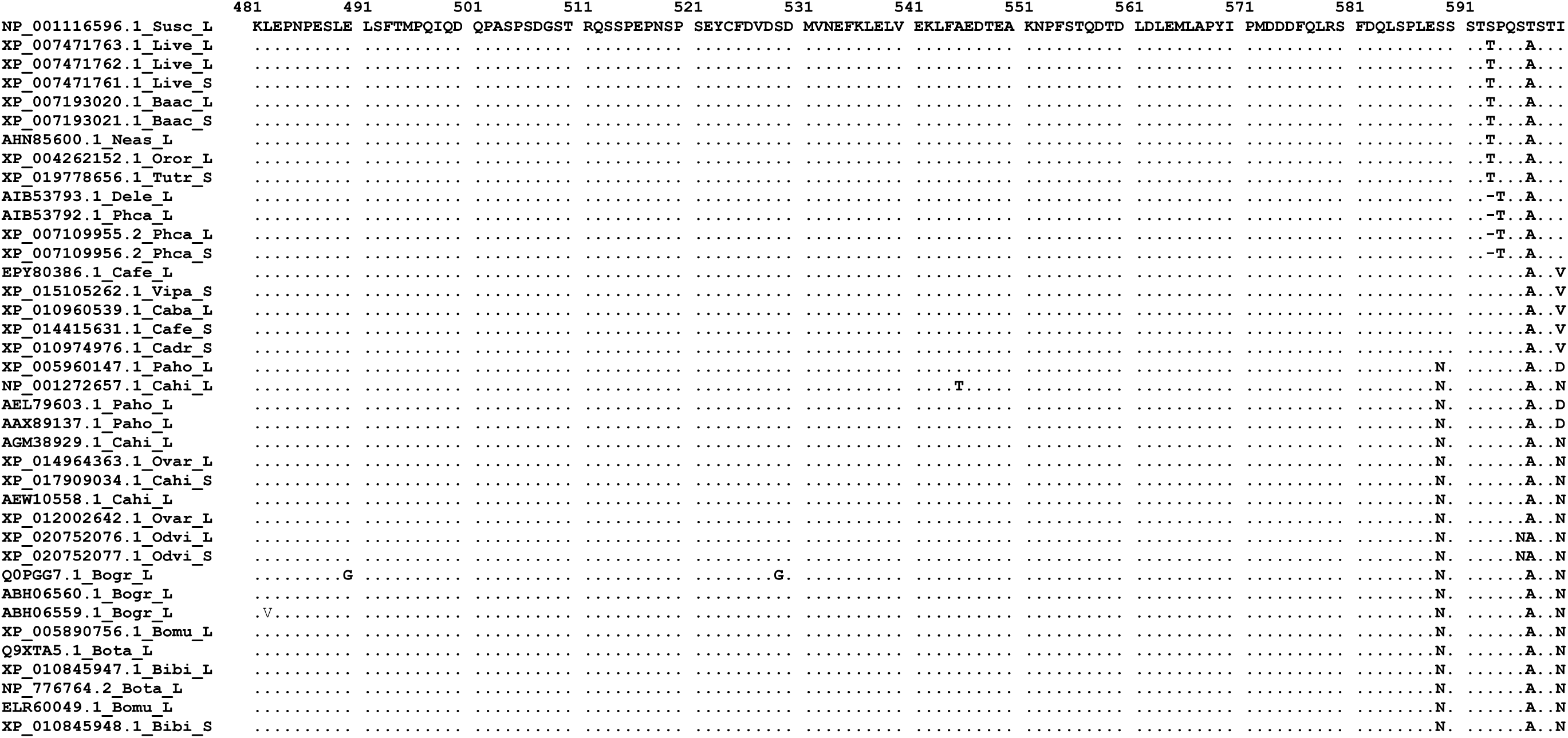
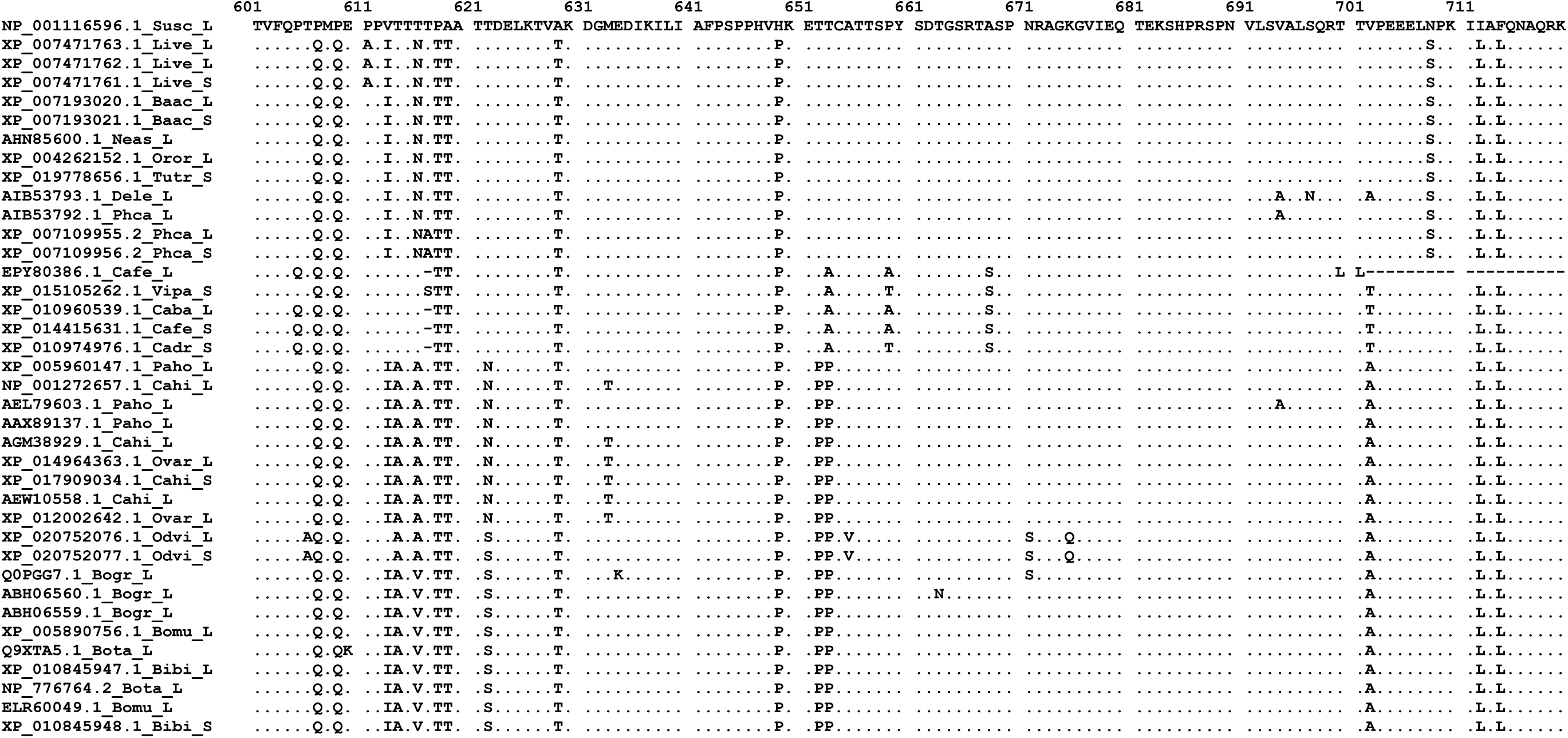
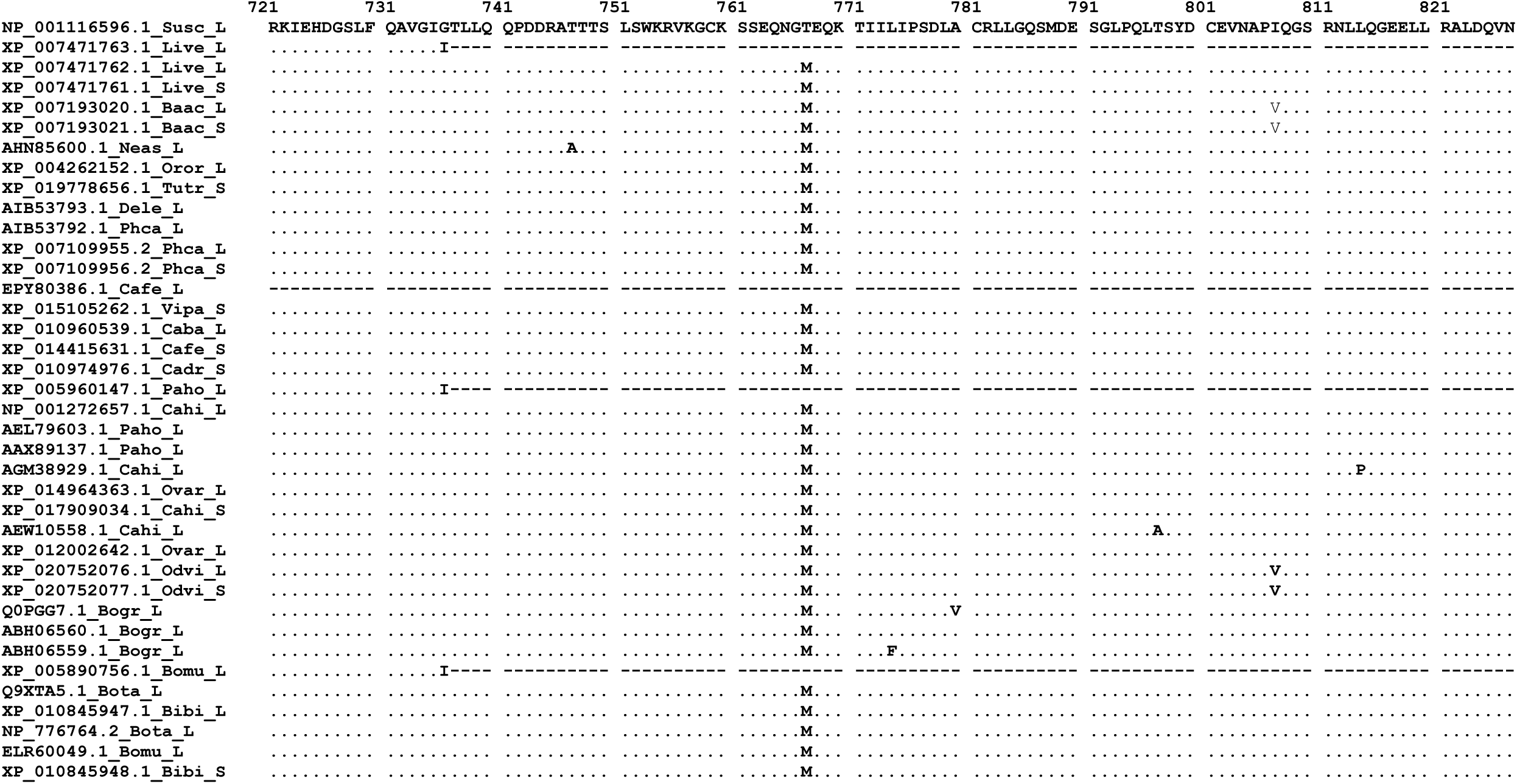

## Supplementary material B

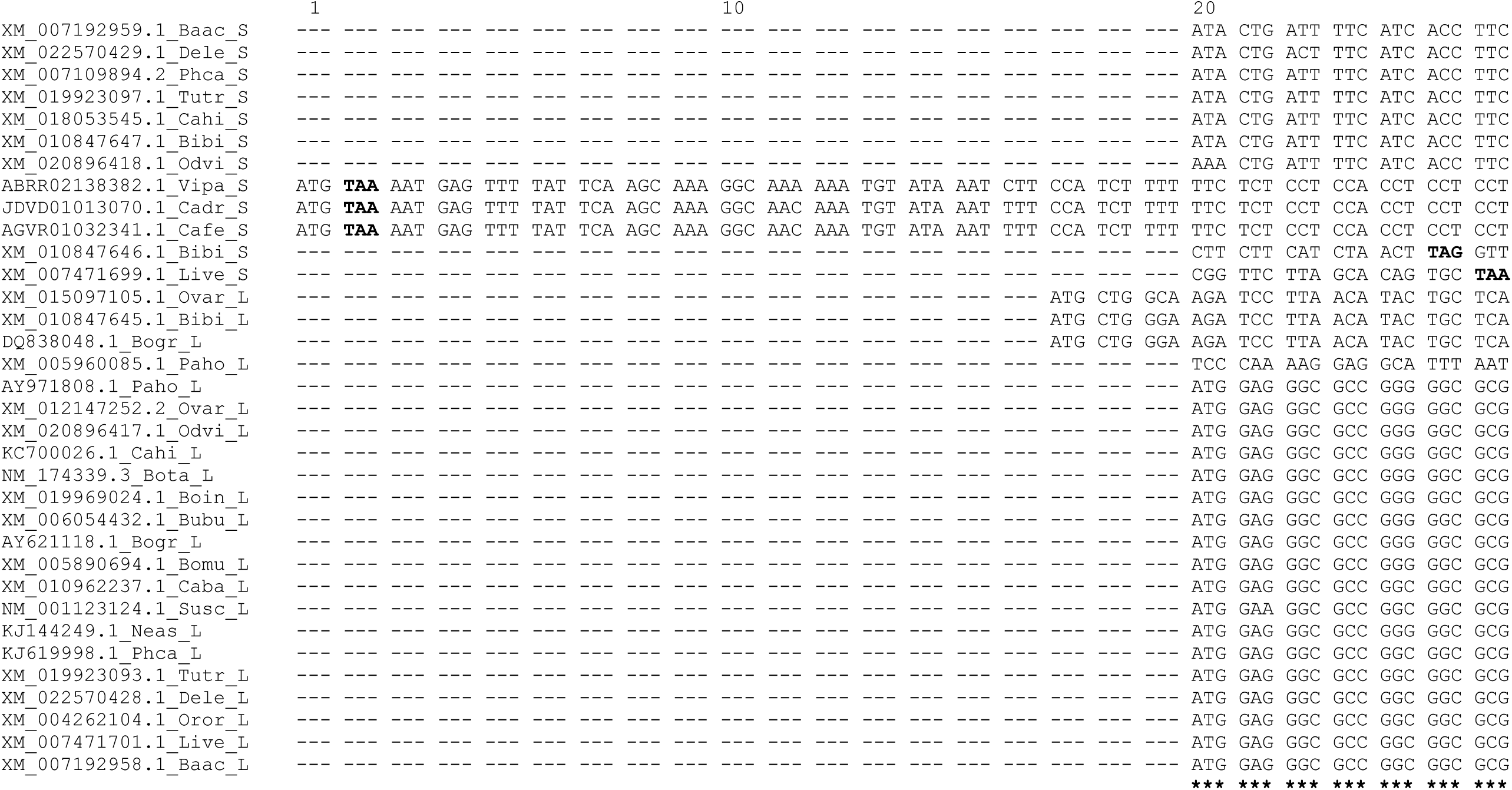
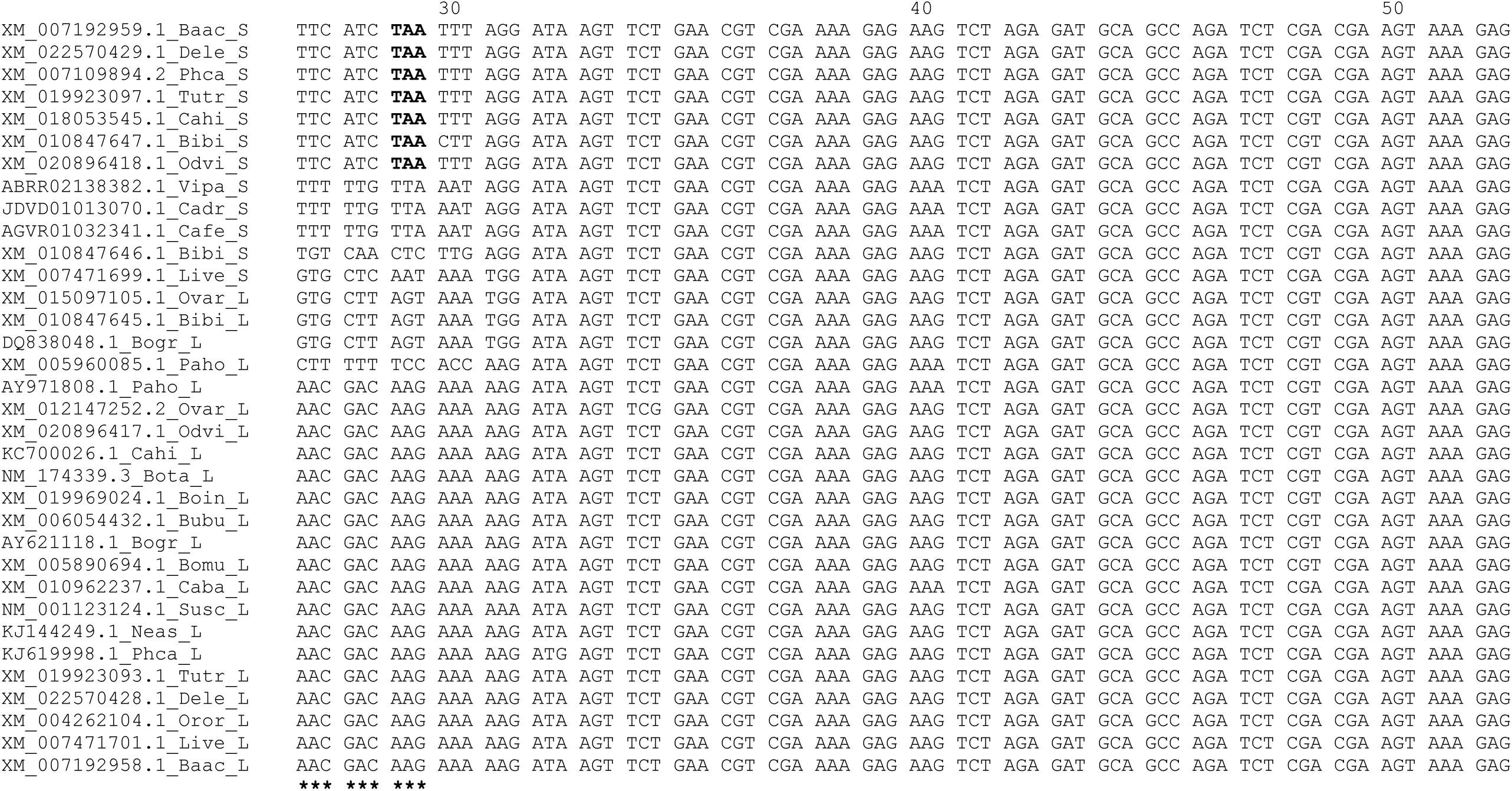
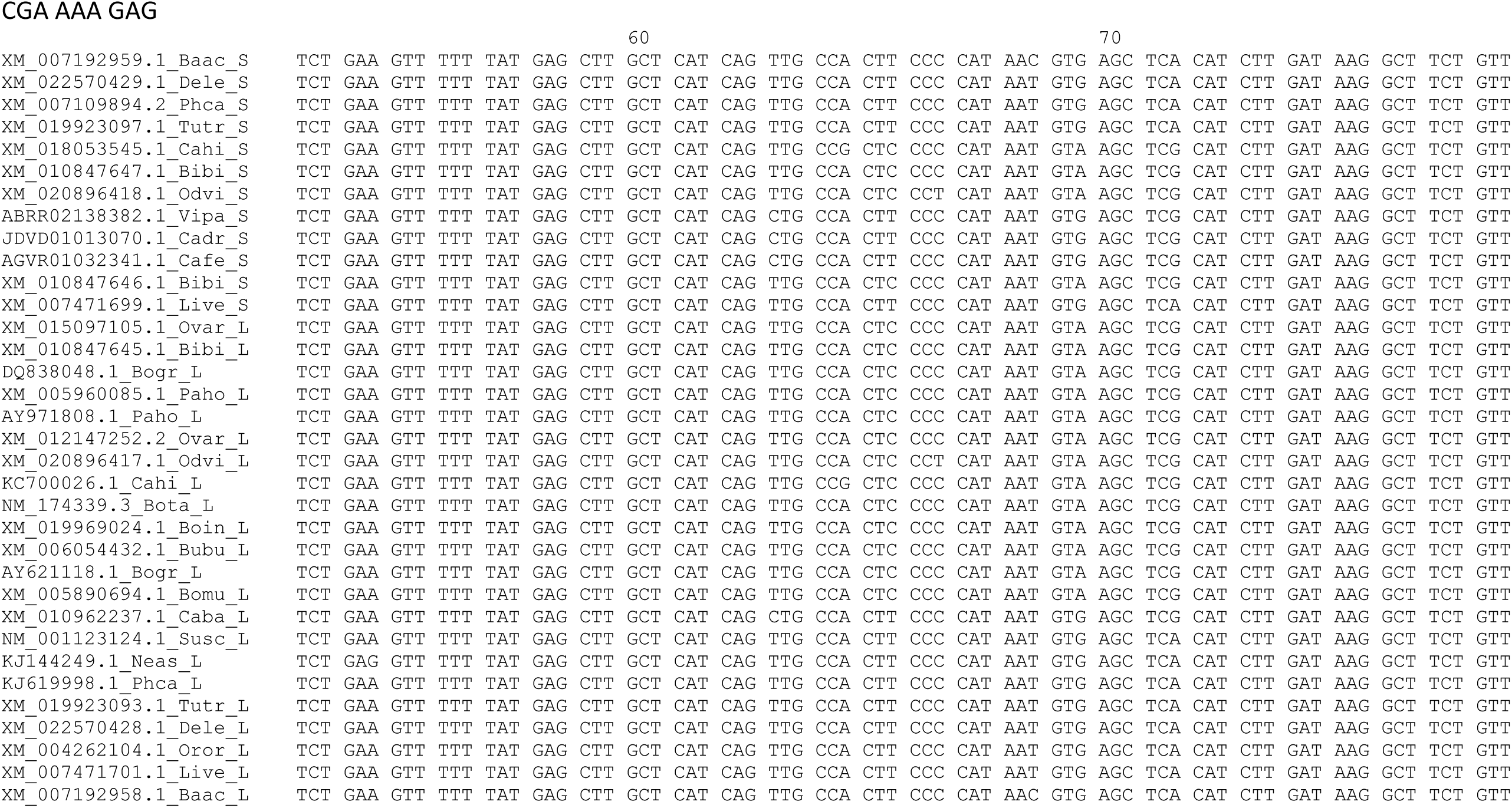

